# Integrated *in silico* and 3D *in vitro* model of macrophage migration in response to physical and chemical factors in the tumor microenvironment

**DOI:** 10.1101/2020.02.17.953208

**Authors:** Sharon Wei Ling Lee, R.J. Seager, Felix Litvak, Fabian Spill, Je Lin Sieow, Hweixian Leong Penny, Dillip Kumar, Alrina Shin Min Tan, Siew Cheng Wong, Giulia Adriani, Muhammad H Zaman, Roger Dale Kamm

## Abstract

Macrophages are abundant in the tumor microenvironment (TME), serving as accomplices to cancer cells for their invasion. Studies have explored the biochemical mechanisms that drive pro-tumor macrophage functions, however the role of TME interstitial flow (IF) is often disregarded. Therefore, we developed a three-dimensional microfluidic-based model with tumor cells and macrophages to study how IF affects macrophage migration and its potential contribution to cancer invasion. The presence of either tumor cells or IF individually increased macrophage migration directedness and speed. Interestingly, there was no additive effect on macrophage migration directedness and speed under the simultaneous presence of tumor cells and IF. Further, we present an *in silico* model that couples chemokine-mediated signaling with mechanosensing networks to explain our *in vitro* observations. The model proposes IL-8, CCL2 and β-integrin as key pathways that commonly regulate various Rho GTPases. In agreement, *in vitro* macrophage migration remained elevated when exposed to a saturating concentration of recombinant IL-8 or CCL2, or to the co-addition of a sub-optimal concentration of both cytokines. Moreover, antibody blockade against IL-8 and/or CCL2 inhibited migration that could be restored by IF, indicating cytokine-independent mechanisms of migration induction. Importantly, we demonstrate the utility of an integrated *in silico* and 3D *in vitro* approach to aid the design of tumor-associated macrophage-based immunotherapeutic strategies.

## INTRODUCTION

Interstitial flow (IF) is an important, yet underappreciated, biophysical force that drives cancer progression [1]. It is derived from an elevated interstitial fluid pressure (IFP) in the solid tumor (∼ 10-40 mmHg) [2, 3], due to highly permeable tumor vessels and the lack of functional lymphatic vessels [4, 5]. This abnormal pressure results in a steep pressure gradient near the tumor margin with escape of interstitial fluid from the tumor mass into the surrounding tissues [6–8], where IFP then drops rapidly to normal tissue values (∼ 0 mmHg) [4, 9]. Because this fluid contains tumor-secreted cytokines and growth factors and can influence tumor cell migration, IF essentially fuels tumor metastasis [9, 10].

Clinical data across different solid cancers have shown that higher IFP at the tumor site strongly correlates with poorer patient survival [11–14]. Accordingly, IFP has been viewed as a strong prognostic factor that is independent of other clinical parameters [11, 15], with ongoing effort to develop strategies to decrease IFP, and hence IF, in patients [16–21]. However, given the complexity of cancer pathology, current findings have only begun to explain the multiple interacting facets of IF’s role in cancer, including its regulation of pivotal immune cell players in the tumor microenvironment (TME). This scenario motivates the need for additional studies that investigate the role of IF on multiple cell types in the TME.

Macrophages are highly abundant at the tumor-stromal boundary [22–24], where there are high levels of IF [25, 26]. Interestingly, this is also where there are high rates of tumor cell invasion [25, 26]. Clinical and experimental evidence report that macrophages crucially support tumor metastasis [27], with a meta-analysis showing that over 80% of studies correlate poor patient outcomes and macrophage density [28]. Moreover, intravital imaging of fluorescently labelled cells in mammary tumors has shown that tumor cells and macrophages move concordantly [24]. Indeed, growing evidence suggests that macrophages could be importantly involved in IF-based tumor cell invasion [25, 29]. Therefore, interfering with the signaling pathways associated with IF, tumor cells and macrophages could potentially inhibit the pro-tumor function of macrophages and also tumor cell metastasis [30].

Macrophages remodel extracellular matrix (ECM) through matrix metalloproteinases (MMP) which degrade collagen and create tracks for tumor cells to migrate [31, 32]. Such tracks also enable macrophages to migrate toward and interact with other cells in the TME to support tumor progression, for example by their contact-dependent support of the epithelial-to-mesenchymal transition (EMT) of tumor cell aggregates [33], or their contact with the endothelium to increase its permeability to intravasating tumor cells [34]. Therefore, macrophage migration in the TME is an important parameter that reflects their ability to support tumor cell invasion through the ECM in the process of metastasis [35]. Current understanding of macrophage migration has been confined to the regulation by biological cues in the TME, including tumor-secreted factors (TSF) which have been identified as key regulators of macrophage motility [36–40].

The study of cell migration requires suitable experimental models that allow cells to migrate with spatial and temporal freedom, while allowing for real-time measurements. A classical set-up is the transwell migration assay, where cell motility is assessed by the number of cells that migrate across a two-dimensional (2D) porous membrane between upper and lower chambers [41, 42]. While the use of such platforms contributes insight toward cell motility, they lack a 3D ECM and fail to accurately mimic the physiological setting [43, 44]. Specifically, recent studies have reported differences in protein expression when cells migrate through a 3D matrix compared to their migration on a 2D substrate [42,44–46]. For example, focal adhesion kinase (FAK) is crucial for migration through a 3D matrix, but in 2D, FAK-null cells can compensate for migration defects by over-expressing other cell migration machineries [46]. Moreover, matrix degradation, an important factor in 3D migration, is not required for migration on a 2D surface. Indeed, there is evidence that 3D measurements of migration directedness (ability to maintain direction of motion) and speed do not correlate with migration in 2D [45]. Finally, these classical assays present end-point readouts of the number of transmigrated cells, failing to capture the temporal dynamics of cell motility.

On the other hand, microfluidic models present suitable 3D environments for real-time measurements of migration directedness and speed. However, current microfluidic models of IF comprise a gel-based mono-culture of cells including tumor cells [47, 48], fibroblasts [49, 50] or macrophages [51]. The most cellularly complex model has tumor cells seeded in a central microchannel with an additional endothelial monolayer in the adjacent microchannel [52]. Recently, Li *et al*. report evidence that IF increased the migration directedness and speed of mouse macrophages using a single-gel microfluidic platform [51]. However, there remains open questions such as how IF which specifically arises from tumor cells can dynamically modulate macrophage migration, and whether IF or TSF is the stronger determinant of macrophage migration behavior. To this end, the *in silico* modelling of *in vitro* data can yield quantitative insight into the biological signalling and biomechanics of macrophage migration in a 3D TME setting [53, 54].

The present work seeks to address these unanswered questions by developing a human-based microfluidic model comprising a co-culture of tumor cells and primary monocyte-derived macrophages, designed to impart the effect of tumor-derived IF and TSF on macrophages. Specifically, we will contribute insight toward the dynamic IF-associated interplay that exists between tumor cells and macrophages. Further, we will clarify if IF and TSF are functionally redundant or additive in regulating macrophage migration, thus identifying their relative importance as potential therapeutic targets. Moreover, we explore how our *in vitro* data contributes toward the development of a refined *in silico* signaling network model that associates TSF, IF and the migration activity of macrophages.

## MATERIALS AND METHODS

### Generation and culture of GFP stable cell lines

A human pancreatic adenocarcinoma (PDAC) cell line, Panc1 (ATCC^®^ CRL-1469), and normal pancreatic epithelial cell line hTERT-HPNE (“HPNE”) (ATCC^®^ CRL-4023) were transfected to stably express green fluorescent protein (GFP). GFP gene was amplified from pGreenPuro shRNA Cloning and Expression Lentivector (CMV; System biosciences, SI505A-1) and sub-cloned into ITR-CAG-DEST-IRES-Neomycin-ITR plasmid (generous gift from Marc Supprian Schmidt). ITR-CAG-DEST-IRES-Neomycin-ITR (control plasmid) (Supplementary Fig. S1a) or ITR-CAG-GFP-IRES-Neomycin-ITR (GFP plasmid) (Supplementary Fig. S1b) was mixed with SB100x transposes (1:1 ratio) and the mixture was transiently transfected into the cell line using Lipofectamine 2000 (Thermo Fisher Scientific, 11668027). Three days post-transfection, transfected cells were selected with 300 µg/mL G418 for 10 days. To validate the stable and constitutive expression of GFP, selected cells were analyzed by flow-cytometry. GFP-expressing Panc1 and HPNE were cultured in Iscove’s Modified Dulbecco’s Media (IMDM; GE Healthcare Hyclone, SH30228.01) supplemented with 5% human serum (Innovative Research, IPLA-SER) and 1% 1X Penicillin-Streptomycin (hereafter referred to as “cIMDM”). Cells were maintained in a humidified CO_2_ incubator at 37 °C and 5% CO_2_.

### Multiplex array

Conditioned media was generated from either the 2D culture of Panc1 and HPNE cells using a previously described method [55] or the 3D culture of these cell lines. Briefly, in the 2D culture, 1 × 10^6^ cells were seeded in 30 mL of cIMDM in a T175 flask and allowed to grow to 70-80% confluency. Media was then removed from the flasks, centrifuged for 10 min at 15000 rpm, sterile-filtered (0.2 µm pore size) and stored at −20 °C until use. For the 3D culture, cells were seeded at equal densities in the microfluidic device for at least 24 h, before collecting media from all media reservoirs of each device. Cell media was centrifuged at 14000 rpm for 10 min at 4 °C and the supernatant was stored in –20 °C until use. 2D and 3D culture media cytokines were respectively analyzed by the Proteome Profiler^™^ antibody array (R&D Systems) and the Milliplex 38 Cytokine kit (Millipore, HCYTMAG-60K-PX38).

### Isolation of monocytes and differentiation into macrophages

Blood samples and procedures used in this study have been approved by the Centralised Institutional Review Board, SingHealth (reference no: 2017/2512) and the Committee on the Use of Humans as Experimental Subjects (COUHES). All protocols are in accordance with The Code of Ethics of the World Medical Association. Written informed consent was given according to the principles expressed in the Declaration of Helsinki. Peripheral blood mononuclear cells (PBMCs) were isolated from whole blood of healthy donors by Ficoll-Paque (GE Healthcare, 17-1440-02) density gradient centrifugation, and monocytes were positively isolated using CD14 microbeads (Miltenyi Biotec, Auburn, CA). Monocytes were maintained in Petri dishes in cIMDM and 100 ng/mL recombinant human M-CSF (Immunotools, Friesoythe, Germany) over 7 days to generate macrophages. Cell viability was assessed by Trypan blue exclusion and was consistently > 90% viable.

### Fabrication of microfluidic device

Microfluidic devices were fabricated following previously reported protocols [56, 57]. Polydimethylsiloxane (PDMS; Sylgard 184 silicone elastomer kit, Dow Corning, Midland, MI, USA) were fabricated by standard soft lithography methods from a patterned SU-8 silicon wafer. Silicone elastomer and curing agent were mixed at a 10:1 weight ratio, degassed in a desiccator, poured onto the photolithographically patterned SU-8 structures and cured overnight at 37 °C. Devices were cut from the PDMS replica, and inlet and outlet ports were created by biopsy punches before autoclave sterilization. After drying the devices overnight at 80 °C, PDMS layers were plasma bonded to the glass cover slips to create channels of approximately 190 μm in height. Each device consists of four connected channels (4.36 mm in length), with two for injecting hydrogels (580 μm wide) and two for culture media (920 μm wide) (Fig. 1a). Each gel channel contains 9 trapezoidal structures (base lengths of 290 μm and 120 μm, height of 140 μm) [33,55,56].

**Figure 1.**
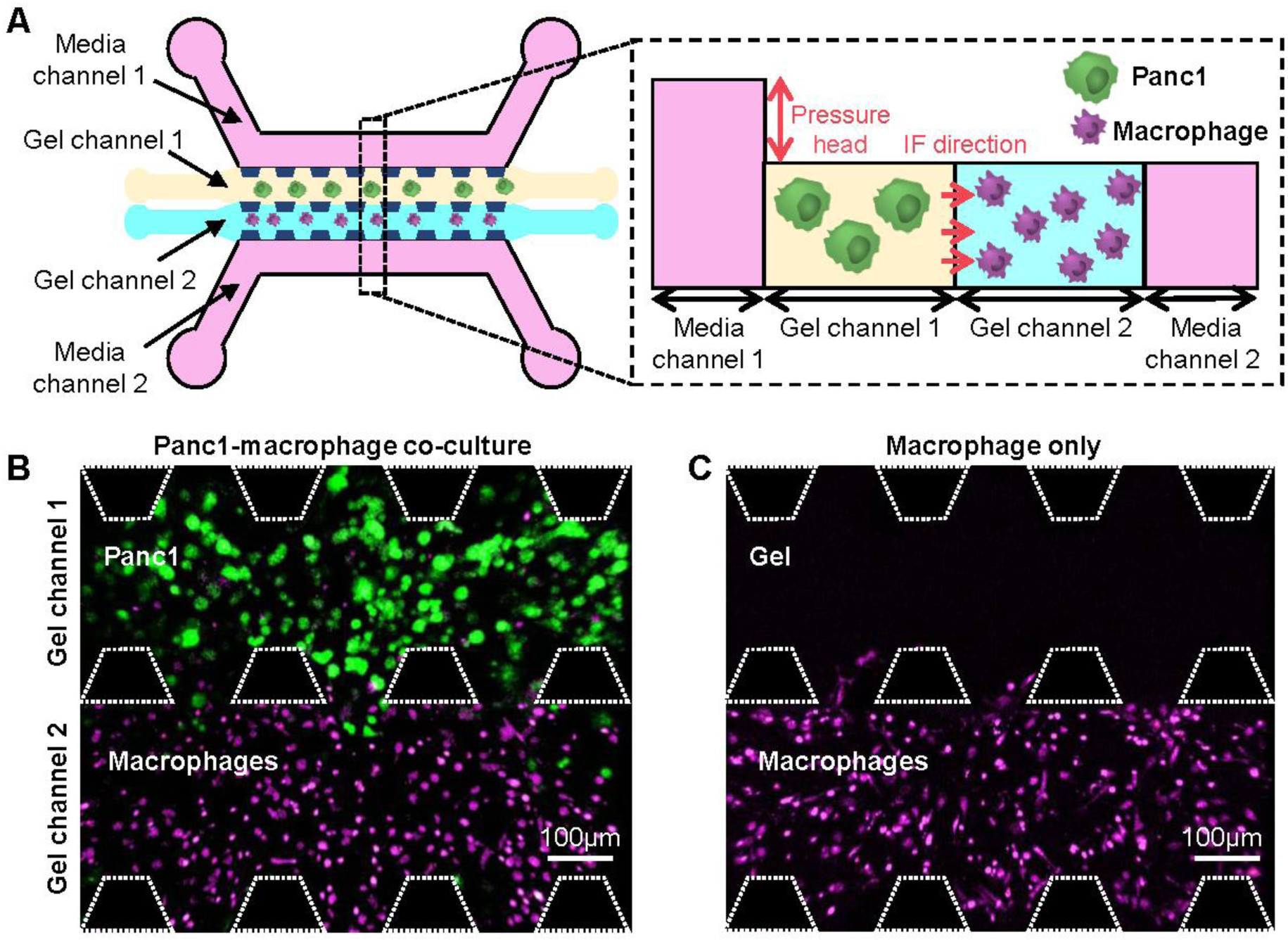
(A) Design of microfluidic co-culture model of tumor cells (Panc1) and macrophages with the incorporation of interstitial flow (IF). Representative confocal image of cell-seeded gel channels at 0 h (B) with tumor cells or (C) without tumor cells.

### Cell seeding

All channels were coated with 1 mg/mL poly-D-lysine (PDL) solution (Sigma-Aldrich, St. Louis, MO) to prevent the detachment of collagen gels from the channel walls [43]. GFP-Panc1 were trypsinized, counted and resuspended in 2.5 mg/mL type I rat tail collagen gel (354236, Corning) solution. Macrophages were harvested by rinsing with D-PBS, incubating with PBS/EDTA (PBS; 2 mM EDTA; Axil Scientific, BUF-1052) for 10–15 min at 37 °C, 5% CO_2_, before adding cIMDM and gently scraping. Harvested macrophages were fluorescently stained with 2 µM Cell Tracker Orange, CMRA (Invitrogen) and resuspended separately in the same type of hydrogel. Cell seeding was performed using a protocol that was previously described [33,55,56]. Briefly, the GFP-Panc1 hydrogel suspension was injected into one gel channel and allowed to polymerize for 20 min in the incubator (37 °C, 5 % CO_2_), followed by injection of the macrophage hydrogel suspension in the other gel channel and then gentle addition of cIMDM into the lateral fluidic channel adjacent to the GFP-Panc1 gel channel. The device was returned to the incubator for 40 min for the second injected hydrogel to polymerize before gentle addition of cIMDM to the media channel adjacent to the macrophage gel channel. In some devices, collagen hydrogel without GFP-Panc1 (herein referred to as “blank hydrogel”) was used as the first hydrogel being injected (Fig. 1b and 1c). Devices containing cells were left to stabilize overnight in the incubator.

### 3D assay and quantification of macrophage migration

After overnight incubation, the cell-containing PDMS chamber was sealed against another PDMS layer that contained a large media reservoir (Fig. 1a). To achieve a flow velocity of ∼ 3 µm/s through the 3D collagen hydrogel, a media-height difference of 2.5 mm was established at the inlet ports connecting the media reservoir with the channels containing GFP-Panc1 or blank hydrogel. Through this set-up, media flowed from the media reservoir, through the GFP-Panc1 cells (or blank hydrogel), and then finally to the macrophages. Darcy’s law was used to calculate the required media-height difference across the gel channels (refer to ‘Calculation and verification of interstitial flow’). In some devices, respective concentrations (ranging from 25 ng/mL to 100 ng/mL) of recombinant human IL-8 (Biolegend, 715404) and/or recombinant human CCL2 (Peprotech, 300-04) reconstituted in cIMDM was added. In other devices, 0.4 µg/mL of anti-IL-8 antibody (R&D, MAB208) and/or 1 µg/mL of anti-CCL2 antibody (R&D, MAB279) was added.

Devices were transferred onto a confocal microscope (Olympus model FV1000) fit with a humidified environmental chamber which maintained a temperature of 37 °C and 5% CO_2_. Macrophages were exposed for 24 h to the various stimuli, including IF, recombinant cytokines and anti-IL-8/anti-CCL2 antibodies, where their migration in 3D was tracked by time-lapse confocal microscopy, with 3D image stacks taken every 25 min at a 20X magnification (800 × 800 pixel density). As the cross-sectional area of the reservoirs was approximately 1500 times that of the hydrogel region, there was a negligible decrease in the media-height difference during the 24 h duration. IMARIS 9.2 was used to track and quantify the migration directedness and speed of macrophages in 3D and to produce the cell trajectory plots of macrophage migration.

### Calculation and verification of interstitial flow

The hydraulic permeability, *K,* of the 2.5 mg/mL collagen gel used in this study was previously determined by Darcy’s law to be ∼ 7 x 10^-14^ m^2^ [51]. Darcy’s law was also used to determine the media-height difference that was required to establish a desirable IF velocity, *v*, through the microfluidic device, as represented by Eq. 1:

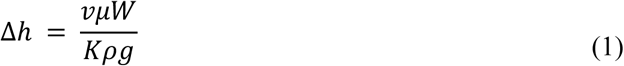

where *ρ* is the density and *μ* is the viscosity of cIMDM media, *W* is the width of the chamber, and Δ*h* is the initial media–height difference of the reservoir [58]. Based on the geometry of the total hydrogel matrix in the microfluidic device and the estimated hydraulic permeability, a media–height difference of 2.5 mm was needed to generate an IF velocity of approximately 3 μm/s in the hydrogel.

To quantify and confirm the velocity of IF through the hydrogel at the start and end of the IF treatment, fluorescence recovery after photobleaching (FRAP) was used. Here, 100 µg/mL of 70 kDa FITC-dextran in cIMDM was added and a spot of 30 µm diameter of the hydrogel was bleached using the highest intensity of the laser, according to the microscope guidelines. After the bleaching step, time-lapse images in short intervals (∼ 1.6 s) were recorded to monitor the recovery in fluorescence. Photobleaching was performed for the hydrogels in both gel microchannels. ImageJ and Matlab was used to quantify the change in fluorescence after the bleaching step.

### Signaling network model construction

A simplified version of the hypothesized signaling network model linking IL-8-based and CCL2-based signaling and IF-induced macrophage migration was constructed by analysing literature-established signaling pathways associated with each of these extracellular stimuli. Relevant networks were analyzed for intersections (which indicate the presence of key signaling species), and common regulators were used to combine and unify the networks into a single network. These common regulators were determined to be various members of Rho GTPases or regulatory signaling species associated with their activity. The similar trends exhibited by both macrophage migration directedness and speed also suggested a common regulator. Thus, we combined the downstream signaling activity of Rho GTPases into a single regulatory node that regulated both migration directedness and speed. Finally, we simplified the network branches by connecting each upstream stimulus with a common downstream migration regulator, removing all intermediate reactions.

### Mathematical model development

A mathematical model was constructed from the aforementioned hypothesized signaling network model. Specifically, an ordinary differential equation (ODE) model tracked the concentration of each signaling element in its active form and its interactions with other signaling elements [59, 60]. The concentrations of activated receptors CXCR1/2 and CCR2 were defined in relation to the concentrations of their respective ligands IL-8 and CCL2 using steady state approximations and the ligand-receptor dissociation constant, *k_d_* (Eq. 2 and 3) [61]. The concentration of activated FAK in response to integrin-mediated, IF-induced signaling was calculated using a Hill function. Here, IF speed was correlated with the steady state concentration of activated FAK, and constants were calculated to reproduce *in vitro* experimental observations (Eq. 4) [51]. Next, the concentration of active G proteins dissociating from G protein-coupled receptors CXCR1/2 and CCR2 in response to IL-8 and CCL2 binding, respectively, was calculated by integrating the two receptor signals in an additive manner through Hill functions with corresponding dissociation constants (*k_d_*) (Eq. 6). Then, we calculated the concentration of active common regulator (Eq. 8). Here, the concentrations of IL-8, FAK, and CCL2 additively contributed to common regulator activation through Hill functions with corresponding dissociation constants (*k_d_*). Finally, migration directedness and speed were modelled as Hill functions that exhibit basal activity when no active common regulator is present and increase asymptotically to a maximum value as the concentration of active common regulator increases (Eq. 9 and 10). Table M1 defines the signaling species being tracked by the model.

The mathematical modeling framework is as follows (Eq. 2 – 10):

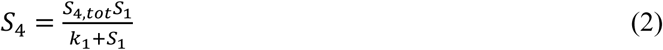

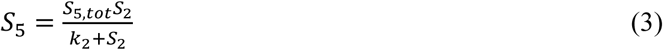

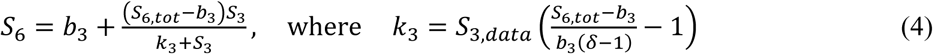

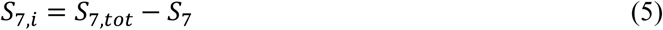

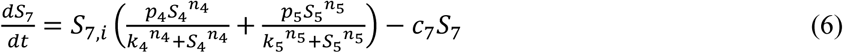

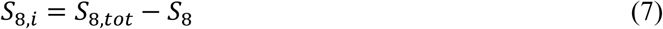

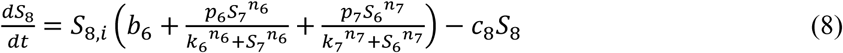

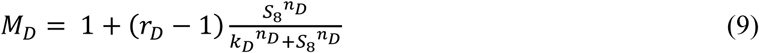

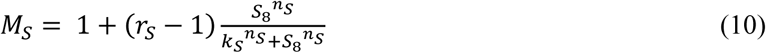

To simulate experiments involving the treatment of macrophages with TSF, or co-culture with cancer cells without IF, the following relationship was used to simulate TSF composition (Eq. 11):

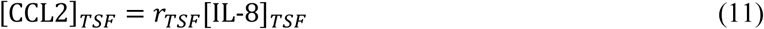

where [IL-8]*_TSF_* and *r_TSF_* are free parameters fit to experimental data.

Furthermore, to simulate experiments where TSF-treated macrophages are exposed to anti-IL-8 and/or anti-CCL2 antibodies, the following functions were used to determine the uninhibited concentrations of each cytokine (Eq. 12 and 13):

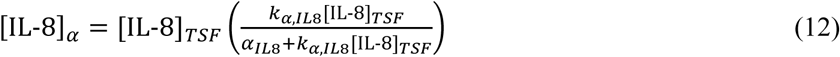

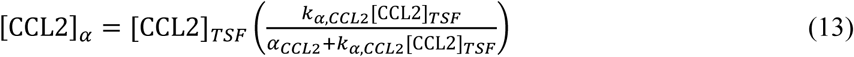

### Model parameter estimation

The mathematical model was implemented as a purpose-constructed code in Python 2.7 and solved at 50,000 time points over 48 h of simulated experimental time *via* the odeint solver found in the scipy.integrate module using the default settings. The model contained 37 parameters, 13 of which were assigned to experimentally-derived values taken from the literature (Table 2), 2 of which were assigned based on manufacturer provided protocols for antibody blockade, and 2 of which were directly calculated from *in vitro* directedness and speed data. Here, the kinetics of each upstream interaction between IL-8, CCL2 or IF speed with downstream signaling elements of the model were taken from the literature. Also, initial conditions of each parameter were determined by running the model without the stimuli until it reached steady state. In addition, the total concentration of each intermediate signaling species (assumed here to be the total concentration of each species at steady state, including both its active and inactive forms) was assigned to a known benchmark value from the literature. The remaining 20 free parameters were fit to our *in vitro* experimental results using a gradient descent least-squares error minimization approach that minimized the error function (Eq. 14):

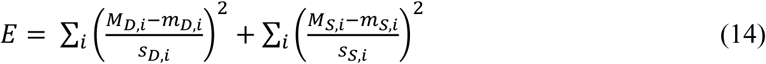

where *M*_*D*,*i*_ and *M*_*S*,*i*_ are the simulated migration directedness and speed for each stimulus, respectively, *m*_*D*,*i*_ and *m*_*S*,*i*_ are the experimentally measured migration directedness and speed for each stimulus, respectively, with associated sample standard deviations *s*_*D*,*i*_ and *s*_*S*,*i*_, respectively. All free parameters were allowed to fit to any value, with the exception of the Hill coefficients (*n_4_*, *n_5_*, *n_6_*, *n_7_*), which were limited to a maximum value of 4. Table 2 shows the fit parameters generated from the model fitting process. Employing such a large number of free parameters proved necessary in order to achieve a sufficiently accurate fit that could capably capture the trends across all 58 experimental data points. We contend that this significant excess in the number of data points — as compared with the number of free parameters — reduced the likelihood of overfitting in this case.

### Sensitivity analysis

Stimulus sensitivity analysis was used to quantify the effect of each extracellular stimulus on the concentration of active common regulator. Here, the elasticity of the concentration of active common regulator was computed for the concentration of each stimulus using the following function (Eq. 15):

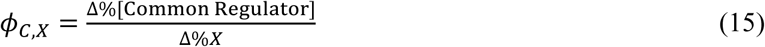

where the denominator represents a 20% change in the stimulus under study (centred at the default stimulus value given in Table 1) and the numerator represents the percent change in the concentration of active common regulator.

**Table 1.** Species variable abbreviations, definitions, initial steady state values and units.

| Species | Abbreviation | Definition | Initial Value | Units |
| --- | --- | --- | --- | --- |
| S<sub>1</sub> | IL-8 | Interleukin-8 | 9.01* | nM |
| S<sub>2</sub> | CCL2 | Chemokine (CC motif) ligand 2 | 9.07* | nM |
| S<sub>3</sub> | IF | Interstitial flow | 3000 | nm/s |
| S<sub>4</sub> | CXCR1/2 | C-X-C motif chemokine receptor 1 or 2; receptor for IL-8 | 0 | nM |
| S<sub>5</sub> | CCR2 | C-C chemokine receptor type 2; receptor for CCL2 | 0 | nM |
| S<sub>6</sub> | FAK | Focal adhesion kinase; activated in response to interstitial flow | 9.48 | nM |
| S<sub>7</sub> | G proteins | Family of signaling G proteins (associated with G-protein-coupled receptors CXCR1/2 and CCR2) | 0 | nM |
| S<sub>8</sub> | Common regulator | Downstream common regulator of migration directedness and speed. Potentially one or more Rho GTPases such as CDC42, Rac1 and/or RhoA | 87.8 | nM |
| M<sub>D</sub> | Directedness | Migration directedness relative to control | 1.00 | - |
| M<sub>S</sub> | Speed | Migration speed relative to control | 1.00 | - |

\*Equivalent to 100 ng/mL.

An additional stimulus sensitivity analysis was used to quantify the effect of each extracellular stimulus on the quality of the model fit against the *in vitro* experimental data. Here, the model *R*^2^ elasticity was computed for each stimulus using the following function (Eq. 16):

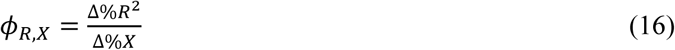

where the denominator represents a 20% change in the stimulus under study (centred at the default stimulus value given in Table 1) and the numerator represents the percent change in the *R*^2^ value between the model outputs and all *in vitro* experimental directedness and speed data.

Parameter sensitivity analysis was used to quantify the effect of each model parameter on the concentration of active common regulator. Here, the elasticity of the concentration of active common regulator was computed for each parameter using the following function (Eq. 17):

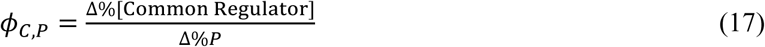

where the denominator represents a 20% change in the parameter under study (centred at the default parameter value given in Table 2) and the numerator represents the percent change in the concentration of active common regulator.

**Table 2.** Parameter values used in simulations. All activation rate constants, dissociation constants, hill coefficients, and basal activation constants are numbered such that they correspond to the reaction numbers indicated in Fig. 4. All total species concentrations and degradation rates are numbered such that they correspond to the species numbers indicated in Fig. 4 and detailed in Table 1.

| Parameter | Value | Units | Definition | Source |
| --- | --- | --- | --- | --- |
| $p_4$ | $9.59 \times 10^{-5}$ | 1/s 1/nM | Activation rate constant for CXCR1/2 activating G proteins | Fit to *in vitro* experimental data |
| $p_5$ | $1.62 \times 10^{-4}$ | 1/s 1/nM | Activation rate constant for CCR2 activating G proteins | Fit to *in vitro* experimental data |
| $p_6$ | $6.97 \times 10^{-3}$ | 1/s 1/nM | Activation rate constant for G proteins activating common regulator | Fit to *in vitro* experimental data |
| $p_7$ | $1.46 \times 10^{-3}$ | 1/s 1/nM | Activation rate constant for active FAK activating common regulator | Fit to *in vitro* experimental data |
| $k_1$ | 1.9 | nM | Dissociation constant for IL-8 binding to CXCR1/2 | [62] |
| $k_2$ | 0.77 | nM | Dissociation constant for CCL2 binding to CCR2 | [63] |
| $k_4$ | 1030 | nM | Apparent dissociation constant for active CXCR1/2 activating G proteins | Fit to *in vitro* experimental data |
| $k_5$ | 739 | nM | Apparent dissociation constant for active CCR2 activating G proteins | Fit to *in vitro* experimental data |
| $k_6$ | 182 | nM | Apparent dissociation constant for G proteins activating common regulator | Fit to *in vitro* experimental data |
| $k_7$ | 183 | nM | Apparent dissociation constant for active FAK activating common regulator | Fit to *in vitro* experimental data |
| $n_4$ | 1.40 | - | Hill coefficient for active CXCR1/2 activating G proteins | Fit to *in vitro* experimental data |
| $n_5$ | 3.42 | - | Hill coefficient for active CCR2 activating G proteins | Fit to *in vitro* experimental data |
| $n_6$ | 4.0 | - | Hill coefficient for G proteins activating common regulator | Fit to *in vitro* experimental data |
| $n_7$ | 1.40 | - | Hill coefficient for active FAK activating common regulator | Fit to *in vitro* experimental data |
| $b_3$ | 9.48 | nM | Basal concentration of active FAK | Fit to *in vitro* experimental data |
| $b_6$ | $1.05 \times 10^{-5}$ | 1/s | Basal rate of common regulator activation | Fit to *in vitro* experimental data |
| $S_{4,tot}$ | 500 | nM | Total concentration of active and inactive CXCR1/2 | [64] |
| $S_{5,tot}$ | 500 | nM | Total concentration of active and inactive CCR2 | [64] |
| $S_{6,tot}$ | 500 | nM | Total concentration of active and inactive FAK | [64] |
| $S_{7,tot}$ | 500 | nM | Total concentration of active and inactive G proteins | [64] |
| $S_{8,tot}$ | 500 | nM | Total concentration of active and inactive common regulator | [64] |
| $c_7$ | $1.56 \times 10^{-4}$ | 1/s | Combined degradation and dilution rate for G proteins | [65] |
| $c_8$ | $1.56 \times 10^{-4}$ | 1/s | Combined degradation and dilution rate for common regulator | [65] |
| $S_{3,data}$ | $3.00 \times 10^3$ | nm/s | Interstitial flow (IF) speed associated with δ-fold increased FAK activation | [48] |
| $\delta$ | 3.28 | - | Fold increase in FAK activation associated with IF at $S_{2,data}$ nm/s | [48] |
| $r_S$ | 3.25 | - | Ratio between highest and lowest observed migration speed | Calculated from *in vitro* experimental data |
| $k_S$ | 202 | nM | Half maximum constant for common regulator-induced migration speed | Fit to *in vitro* experimental data |
| $n_S$ | 3.87 | - | Hill coefficient for common regulator-induced migration speed | Fit to *in vitro* experimental data |
| $r_D$ | 2.95 | - | Ratio between highest and lowest observed migration directedness | Calculated from *in vitro* experimental data |
| $k_D$ | 120 | nM | Half maximum constant for common regulator-induced migration directedness | Fit to *in vitro* experimental data |
| $n_D$ | 3.64 | - | Hill coefficient for common regulator-induced migration directedness | Fit to *in vitro* experimental data |
| $[IL-8]_{TSF}$ | 0.167 | nM | - | Fit to *in vitro* experimental data |
| $r_{TSF}$ | 1.18 | - | - | Fit to *in vitro* experimental data |
| $\alpha_{IL8}$ | 2.67 | nM | Concentration of anti-IL-8 antibody in antibody blockade experiments | Reflective of *in vitro* experimental conditions |
| $k_{\alpha,IL8}$ | 0.888 | - | Molar ratio of antibody to IL-8 required for 50% inhibition | Based on manufacturer supplied datasheet |
| $\alpha_{CCL2}$ | 6.67 | nM | Concentration of anti-CCL2 antibody in antibody blockade experiments | Reflective of *in vitro* experimental conditions |
| $k_{\alpha,CCL2}$ | 1.32 | - | Molar ratio of antibody to CCL2 required for 50% inhibition | Based on manufacturer supplied datasheet |

Finally, an additional parameter sensitivity analysis was used to quantify the effect of each model parameter on the quality of the model fit against *in vitro* experimental data. Here, the model *R*^2^ elasticity was computed for each parameter using the following function (Eq. 18):

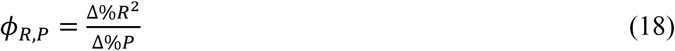

where the denominator represents a 20% change in the parameter under study (centred at the default parameter value given in Table 2), and the numerator represents the percent change in the *R*^2^ value between the model outputs and all *in vitro* experimental directedness and speed data.

### Statistical analysis

Statistical analysis of experimental data was performed using GraphPad Prism 8.2 (GraphPad Software) considering at least two regions of interest (ROI) in the hydrogel per device. For each *in vitro* experiment, the average migration directedness and speed of macrophages in each ROI (∼ 150 macrophages per ROI) was used to generate a data point. To quantify the agreement between the *in vitro* experimental data and the output values of the mathematical model, the *R*^2^ values were calculated for (1) directedness data only, (2) speed data only, and (3) all data together. At each data point, statistical analysis was performed to determine if the population mean predicted by the model would be likely to produce the *in vitro* experimental data. Data were plotted as the mean ± standard error of the mean (SEM), where *n.s.* represents not significant, * represents *P* ≤ 0.05, ** represents *P* ≤ 0.01, *** represents *P* ≤ 0.001 and **** represents *P* ≤ 0.0001. Statistical significance was determined using a Student’s *t*-test or where appropriate, a one-way ANOVA with Holm-Sidak’s multiple comparisons test. Only a *P*-value or adjusted *P*-value of ≤ 0.05 was taken as evidence of statistical significance.

## RESULTS

### Generation of an *in vitro* 3D co-culture model with tumor interstitial flow

Based on previously optimized protocols [51, 56], we designed a 3D *in vitro* microfluidic model of tumor IF to specifically study macrophage migration. In this platform, we cultured tumor cells in one channel and macrophages in the adjacent channel, thus allowing us to simulate and study the impact of tumor-originating IF on macrophage migration without the influence of physical cell contact (Fig. 1). In our set-up, we confirmed that the velocity of IF through both gel channels was within the range reported for tumor tissues [25, 26]. Fluorescence recovery after photobleaching (FRAP) analysis revealed a mean IF velocity of 3.9 ± 0.7 µm/s at the beginning of IF exposure and 4.4 ± 0.5 µm/s after 24 h (*P* > 0.05) (Supplementary Fig. S3). These results correspond with the height difference of 2.5 mm between the two media channels which stayed relatively unchanged over the 24 h. Thus, it could be assumed that IF velocity was relatively constant throughout the duration of IF treatment.

### Similar effect on macrophage migration by tumor-secreted factors and interstitial flow

In our first set of *in vitro* experiments, we observed that compared to the control macrophage monoculture without IF (Fig. 2ai), 24 h of IF exposure appeared to increase macrophage motility (Fig. 2aii) as shown by the relatively increased spread of their x-y migration path trajectories. Interestingly, it appeared that the presence of tumor cells alone without IF (TSF) also increased the spread of migration path trajectories relatively (Fig. 2aiii), suggesting that either IF or TSF alone could promote increased macrophage motility. Interestingly, combining IF and TSF (at the cell concentrations tested in Fig. 2) did not have an additive effect on the spread of macrophage migration (Fig. 2aiv).

**Figure 2.**
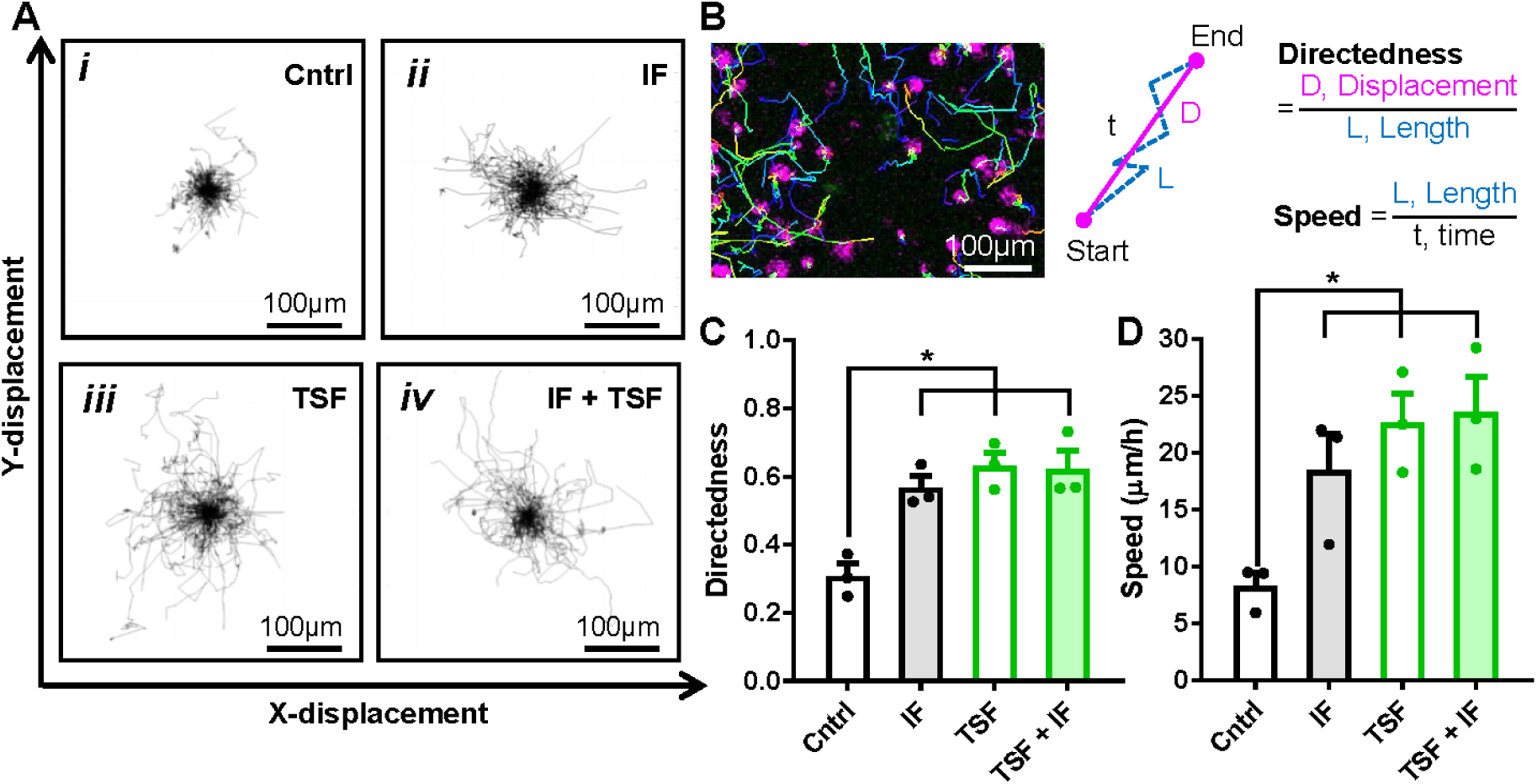
(A) X-Y path trajectories of macrophage migration in the (i) control (Cntrl) monoculture of macrophages without interstitial flow (IF) or exposed to (ii) only IF, (iii) only tumor cells; tumor-secreted factors (TSF), or (iv) both IF and TSF. (B) Quantification method of macrophage migration directedness and speed. (C) Directedness and (D) speed of macrophage migration under the different conditions tested. Data are shown as the mean ± SEM (n = 3), where statistical significance was determined using a one-way ANOVA with Holm-Sidak’s multiple comparisons test with * *P* ≤ 0.05.

To ascertain the effect of each stimulus on macrophage motility, we quantified the directedness and speed of macrophage migration during the period of IF exposure following a previously reported approach (Fig. 2b) [36, 51]. Consistent with the observed increase in x-y migration path trajectories, IF, TSF or the combination of both could substantially increase the directedness (Fig. 2c) and speed of macrophage migration (Fig. 2d). Compared to the untreated control where the directedness and speed of macrophage migration was D = 0.31 ± 0.04 and S = 8.3 ± 1.2 µm/h, respectively, a 2-fold increase in directedness and 3-fold increase in speed was observed for the conditions of IF (D = 0.57 ± 0.03, S = 19 ± 3 µm/h), TSF (D = 0.63 ± 0.04, S = 23 ± 3 µm/h) and the IF-TSF combination (D = 0.62 ± 0.06, S = 24 ± 3 µm/h), and these values are within the range reported in previous *in vitro* investigations [51].

### Hypothesized signaling pathway linking key tumor-secreted factors and Rho GTPase-regulated migration

A multiplex cytokine array of culture supernatant of the tumor cell line (Panc1) and a normal control cell line (HPNE) was conducted to identify the main cytokines within TSF that drove the migration behaviour that we observed *in vitro* of macrophages. Our analysis on 2D cell culture-derived supernatant revealed several cytokines secreted at higher levels by the Panc1 compared to the HPNE cell line (Fig. 3a). A review of known signaling networks revealed that the upregulation of CCL2 and IL-8 is highly associated with the regulation of cell migration [66, 67]. We also confirmed that Panc1 secreted higher levels of both cytokines than HPNE in the 3D *in vitro* culture environment (Fig. 3b), suggesting that IL-8 and CCL2 could mainly drive the migration we observed of macrophages in the 3D *in vitro* system.

**Figure 3.**
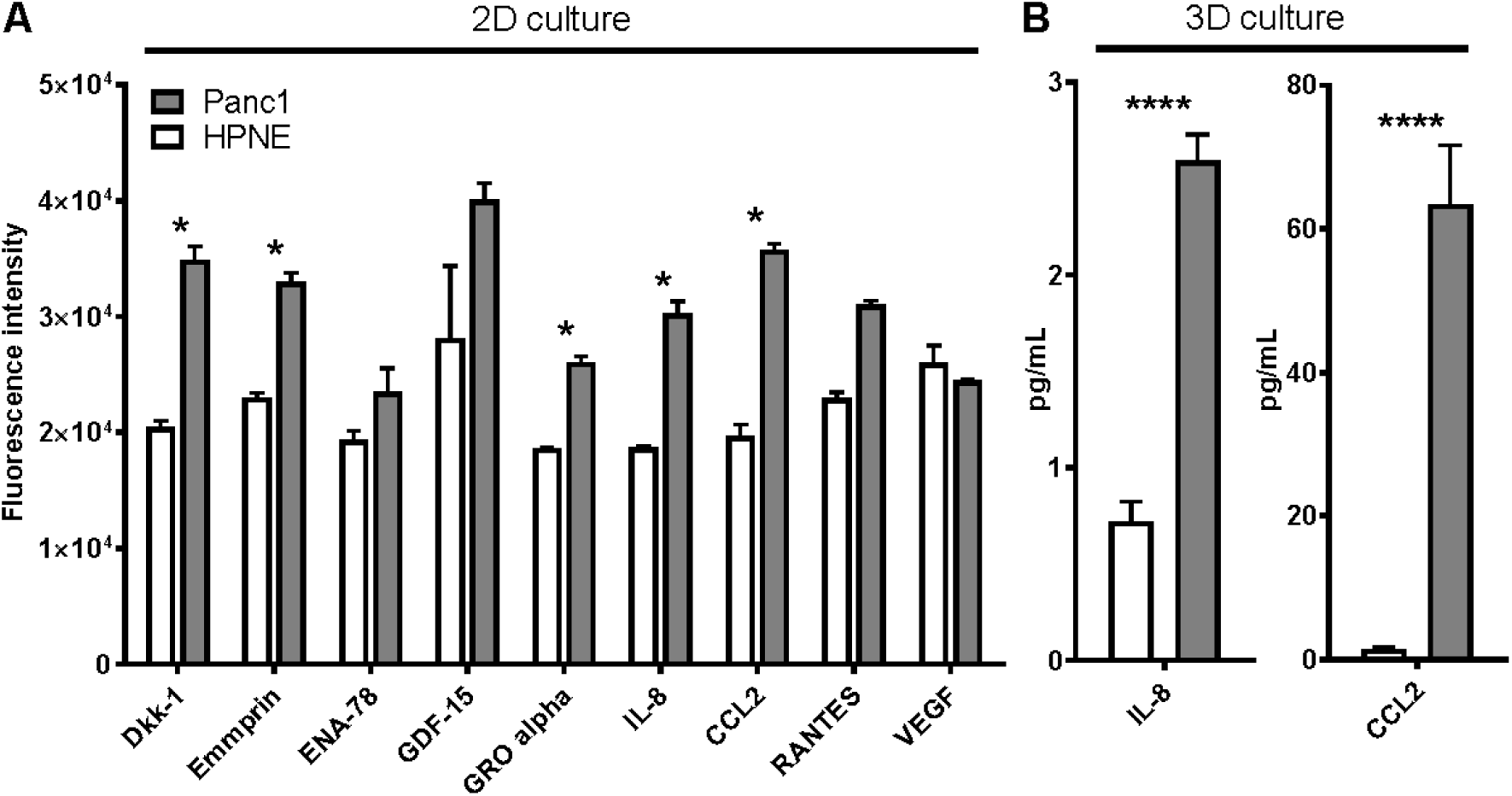
Multiplex array of cytokines in culture supernatant derived (A) from cells grown as a 2D monolayer using the Proteome Profiler^™^ antibody array (R&D Systems) or (B) from cells grown in a 3D matrix environment using the Milliplex 38 Cytokine kit (Millipore). Data are shown as the mean ± SEM (n = 3), where statistical significance was determined using a Student’s *t*-test with * *P* ≤ 0.05 and **** *P* ≤ 0.0001.

Mechanistically, IL-8 [66,68,69] and CCL2 [67,70–73] bind to G protein-coupled receptors CXCR1/2 and CCR2, respectively, resulting in the activation and subsequent dissociation of an associated G protein [74]. This releases the α subunit of the G protein to activate further intracellular signaling that results in the post-translational regulation of regulatory proteins such as small monomeric GTPases of the Rho-family, leading to polymerization and retraction of the actin cytoskeleton which are important processes for cells to migrate. Additionally, our *in vitro* experimental data support previous studies that have demonstrated a synergistic integration between these receptors, whereby downstream signals are significantly greater in response to the activation of both receptors than either receptor alone [75]. This synergy has been shown to depend on the activation of both receptors, suggesting that synergy is the result of intracellular signaling, as opposed to extracellular cytokine interactions or receptor-receptor associations [75]. Moreover, even in the presence of both receptors, certain cell types fail to demonstrate this synergy, suggesting this behaviour arises from a characteristic intracellular signaling motif or protein that is unique to certain cell types such as macrophages [76]. G proteins represent a common element in the downstream signaling networks associated with both receptors [68, 73], and are upstream of migration-regulating Rho GTPases and extracellular signal-regulated kinases (ERK) (which also exhibit a synergistic activation in response to the activation of both receptors) [75]. Thus, we hypothesized that a subset of G proteins may represent the integration point and the source of the synergy between these signaling pathways. In addition, IF-modulated signaling, as mediated by integrin-β2 and FAK, is known to drive the activation and downstream migratory activity associated with Rho GTPases [51, 77]. Finally, because our *in vitro* experimental data show that IF and TSF induced a similar increase in directedness and speed, we hypothesized that IL-8, CCL2 and IF regulate one or more of the relevant Rho GTPases, namely CDC42, Rac1 and/or RhoA [38–40].

Notably, the purpose of our model is to infer the logical structure of the network that integrates multiple molecular (IL-8 and CCL2) and mechanical (IF) stimuli, but not the intricate details of interactions associated with the downstream Rho GTPases. For this reason, we model the Rho GTPases collectively through a single representative concentration, and hereafter denote this group of signaling species collectively as “common regulator”. Also, as downstream signaling (connecting Rho GTPase activity to migration-related processes) becomes increasingly complex, we recognize that there is no need to model all theoretical details of GTPases signaling. Moreover, the similar trends exhibited by both directedness and speed in response to all tested stimuli could be explained simply by a common regulator, as opposed to more complex interactions between largely independent signaling pathways. Thus, we depicted directedness and speed as phenomena that are indirectly induced by this common regulator, with intermediate signaling described by respective response functions that depend on the concentration of active common regulator. These considerations led to our proposed signaling network model (Fig. 4) [37].

**Figure 4.**
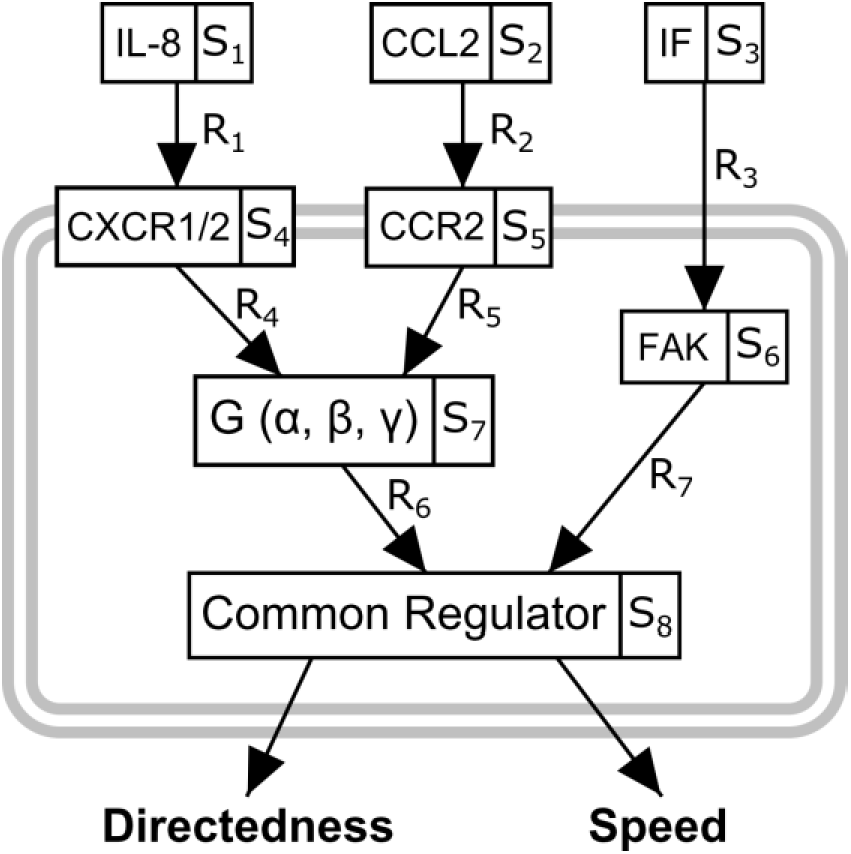
Hypothesized signaling model showing how signals associated with IL-8, CCL2 and interstitial flow (IF) activate a common regulator, both individually and in combination, to regulate macrophage migration directedness and speed. The input stimuli (IL-8, CCL2 and IF), associated cytokine receptors (CXCR1/2 and CCR2) and intermediary signaling species, including focal adhesion kinase (FAK) and heterotrimeric small G-proteins, G(α,β,γ), are identified by their respective abbreviations and species variables (Eq. 2-10 in section ‘Mathematical model development’).

### Effect of varying IL-8, CCL2 and/or interstitial flow on macrophage migration

A second set of *in vitro* experiments were conducted to confirm the central roles of IL-8 and CCL2, and to also validate the proposed signaling network model. Compared to the non-treated control, where macrophage migration directedness and speed were D = 0.35 ± 0.06 and S = 8.4 ± 1.1 µm/h, respectively, the exogenous addition of saturating concentrations (100 ng/mL) of only IL-8 (D = 0.56 ± 0.03, S = 20 ± 2 µm/h) or only CCL2 (D = 0.60 ± 0.07, S = 22 ± 4 µm/h) to a macrophage monoculture substantially increased migration directedness and speed (Fig. 5a). These increases were comparable to those obtained by exposing the macrophage monoculture to only IF (D = 0.61 ± 0.05, S = 23 ± 4 µm/h). Moreover, simultaneously exposing macrophages to IF together with either IL-8 (D = 0.59 ± 0.01, S = 19 ± 4 µm/h) or CCL2 (D = 0.60 ± 0.08, S = 22 ± 3 µm/h) did not further increase macrophage migration (Fig. 5a), supporting the notion that IF and TSF can commonly regulate macrophage migration behaviour.

**Figure 5.**
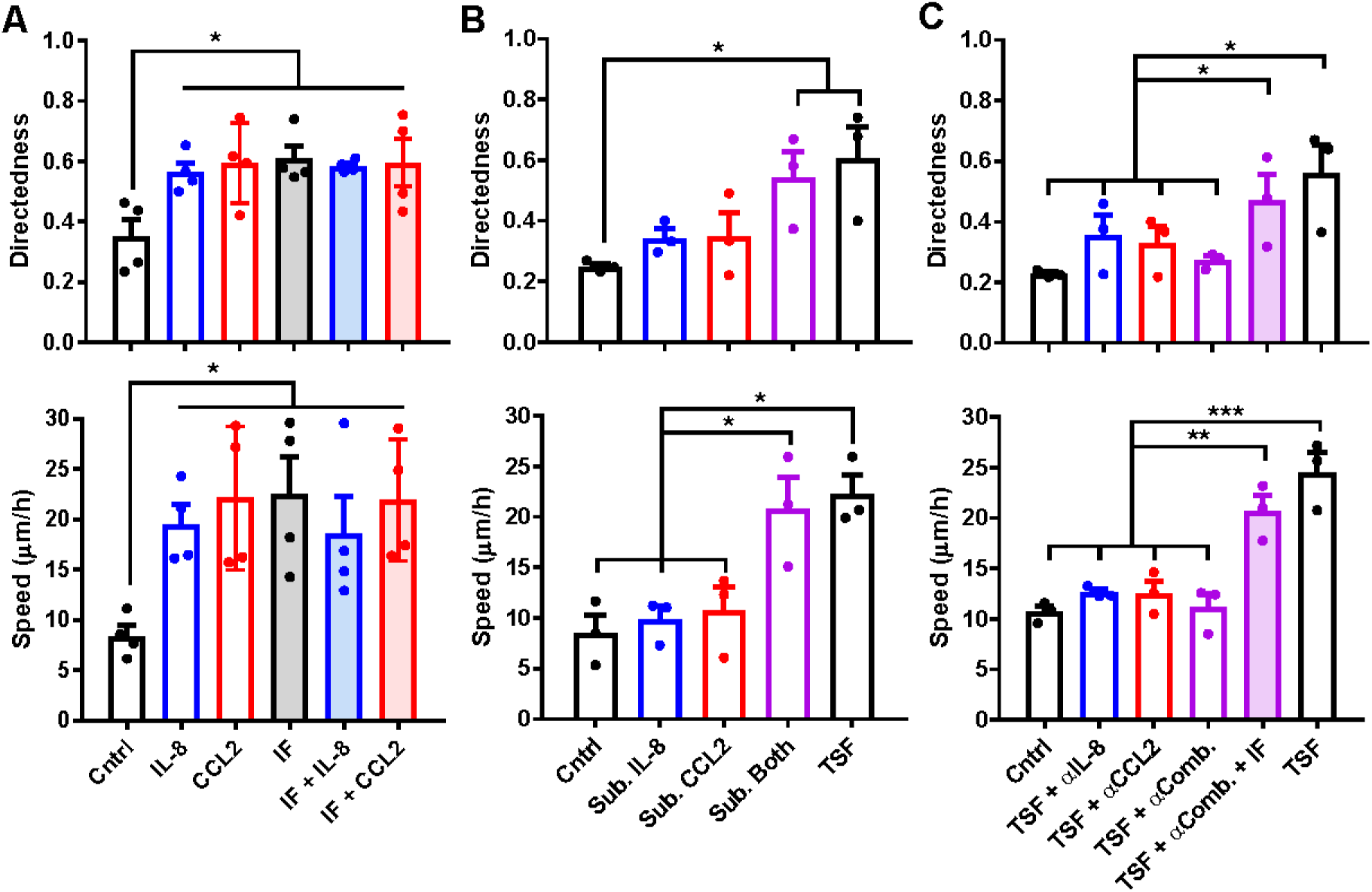
Directedness (top panel) and speed (bottom panel) of macrophage migration with exogenous addition of a (A) saturating concentration (100 ng/mL) of IL-8 or CCL2, or (B) sub-optimal concentration (25 ng/mL) of IL-8 and/or CCL2 to a macrophage monoculture, or the addition of (C) blocking antibodies against IL-8 and/or CCL2 to a macrophage-tumor cell co-culture in the presence or absence of IF. Data are shown as the mean ± SEM (n ≥ 3), where statistical significance was determined using a one-way ANOVA with Holm-Sidak’s multiple comparisons test with * *P* ≤ 0.05, ** *P* ≤ 0.01, and *** *P* ≤ 0.001. (*Cntrl*: control, *IF*: interstitial flow, *Sub.*: sub-optimal, *Sub. Both*: combined addition of sub-optimal concentrations of IL-8 and CCL2, *TSF*: tumor-secreted factors, *αComb.*: combined blockade of IL-8 and CCL2)

Next, macrophages were exposed to sub-optimal concentrations of IL-8 and/or CCL2 to test if the same extent of increase in directedness and speed would be achieved as seen with either (1) a saturating concentration of individual cytokines, or (2) the 3D co-culture of macrophages and tumor cells (TSF). Sub-optimal concentrations of 25 ng/mL of IL-8 and 25 ng/mL of CCL2 were identified based on prior titration experiments (Supplementary Fig. S2a and S2b). While exposure to the sub-optimal concentration of only IL-8 (D = 0.34 ± 0.03, S = 9.9 ± 1.3 µm/h) or only CCL2 (D = 0.35 ± 0.08, S = 11 ± 2 µm/h) did not result in an observable increase in directedness and speed, the combined exposure to sub-optimal concentrations of both cytokines promoted a comparable increase in directedness and speed (D = 0.54 ± 0.09, S = 21 ± 3 µm/h) to the TSF condition (D = 0.6 ± 0.1, S = 22 ± 2 µm/h) (Fig. 5b). Importantly, the data suggest a synergistic integration of the intracellular signals associated with these two cytokines.

Finally, the introduction of 0.4 µg/mL of anti-IL-8 and 1 µg/mL of anti-CCL2 blocking antibodies could inhibit the TSF-mediated increase in migration directedness and speed (Fig. 5c). Compared to the co-culture condition (TSF) (D = 0.56 ± 0.10, S = 25 ± 2 µm/h), both directedness and speed were substantially decreased upon treatment with anti-IL-8 (D = 0.35 ± 0.07, S = 13.0 ± 0.3 µm/h), anti-CCL2 (D = 0.33 ± 0.06, S = 13 ± 1 µm/h) or combinational blockade of both cytokines (D = 0.27 ± 0.02, S = 11 ± 1 µm/h). Interestingly, IF could restore migration back to a level that was comparable to the TSF condition (D = 0.47 ± 0.09, S = 21 ± 2 µm/h) despite combinational antibody blockade of IL-8 and CCL2.

### Modelled signaling network reproduces fundamental migration behaviors

A mathematical model was developed based on the hypothesized signaling network (Fig. 4) using well-established systems biology modeling techniques, in particular the use of Hill functions (1) to relate the concentration of active signaling proteins with the rate of activation of their downstream targets, and (2) to determine the steady state concentrations of bound, active cytokine receptors using dissociation constants from the literature [59, 60]. Here, an ODE model tracked the concentration of each signaling element in its active form and its interactions with other signaling elements. The system successfully replicated the experimental trends observed in Fig. 5a of migration directedness and speed (Fig. 6a). Importantly, the addition of individual or combined stimuli increased directedness and speed to approximately its maximum value, similar to *in vitro* observations. Also, the mathematical model could replicate *in vitro* experimentally observed migration responses to sub-optimal concentrations of IL-8 and/or CCL2 (Fig. 6b), to the antibody blockade of IL-8 and/or CCL2 when tumor cells were present (Fig. 6c), and to the titration of either cytokine concentration in the absence of IF (Supplementary Fig. S2c and S2d). Quantifying model agreement across all data points yields a mean coefficient of determination (*R*^2^) value of 0.70 for all directedness data, 0.90 for all speed data, and 0.83 for all directedness and speed data taken together. Therefore, our model could successfully reproduce 83% of the variation in the *in vitro* experimental data. Additionally, statistical analysis at each data point confirmed a pointwise agreement between *in vitro* experimentally observed and model-generated data, with no substantial difference between the two data types.

**Figure 6.**
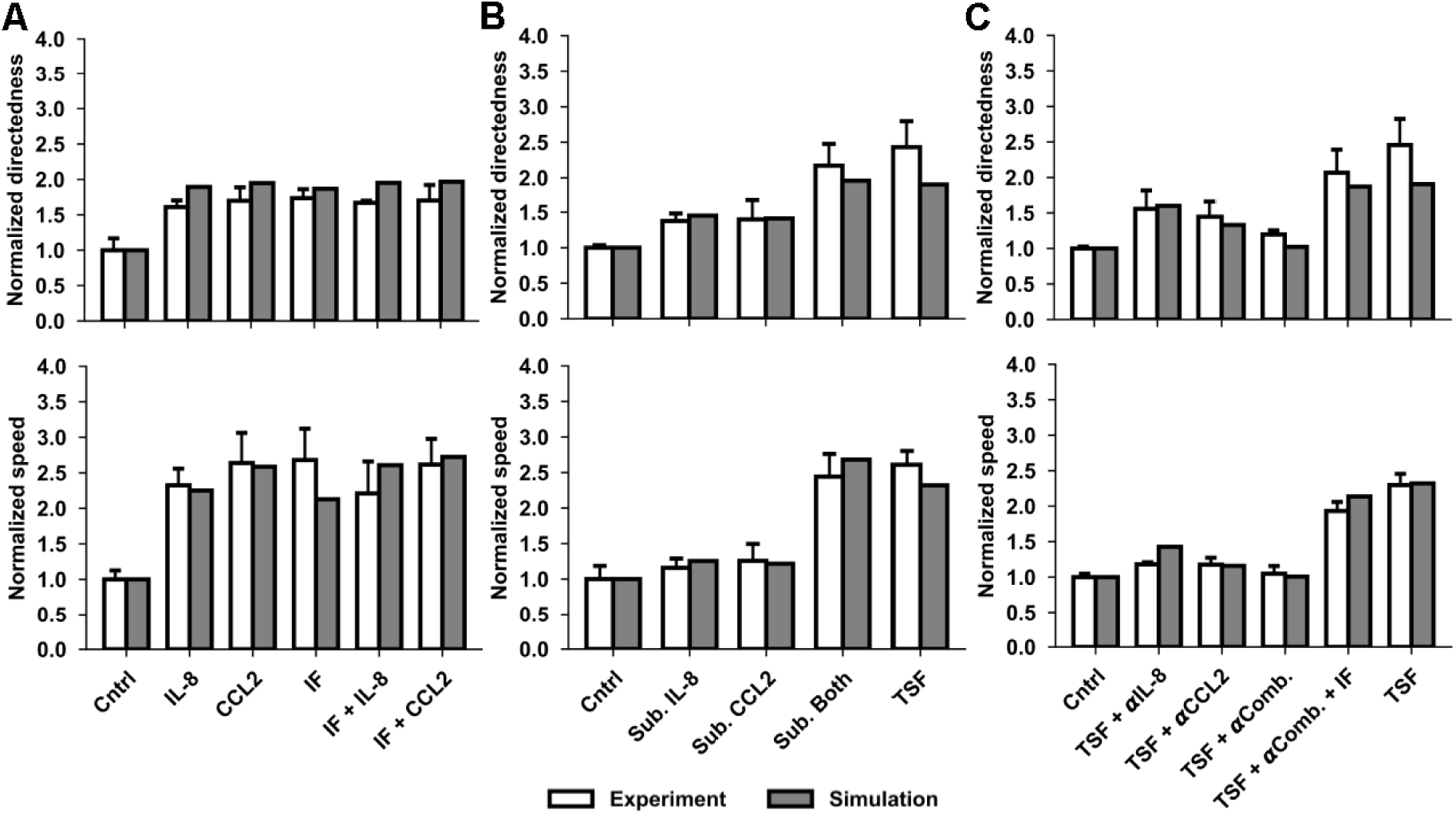
Non-significant difference between *in vitro* experimental (white) and model (grey) predictions for normalized directedness (top panel) and speed (bottom panel) of macrophage migration with exogenous addition of a (A) saturating concentration (100 ng/mL) of IL-8 or CCL2, or (B) sub-optimal concentration (25 ng/mL) of IL-8 and/or CCL2 to a macrophage monoculture, or the addition of (C) blocking antibodies against IL-8 and/or CCL2 to a macrophage-tumor cell co-culture in the presence or absence of IF (100% antibody blockade of cytokines assumed). Data are shown as the mean ± SEM (n ≥ 3), where statistical significance was determined using Student’s *t*-tests which compared between *in vitro* experimental data and model-generated data at a pointwise level with *P* ≤ 0.05. (*Cntrl*: control, *IF*: interstitial flow, *Sub.*: sub-optimal, *Sub. Both*: combined addition of sub-optimal concentrations of IL-8 and CCL2, *TSF*: tumor-secreted factors, *αComb.*: combined blockade of IL-8 and CCL2)

### Sensitivity analysis reveals most influential stimuli and parameters

Sensitivity analyses were conducted to identify the most influential stimuli and parameters that impacted macrophage migration behaviour. Because directedness and speed were both regulated by the common regulator, these analyses captured the influence of each parameter on both types of migration behaviors. First, we performed a stimulus sensitivity analysis to quantify the relative influence of each modelled stimulus on the concentration of active common regulator (Eq. 15) (Fig. 7a). IF had the greatest influence on the concentration of active common regulator, with approximately 1.5 times the influence of IL-8 and over 3 times that of CCL2.

**Figure 7.**
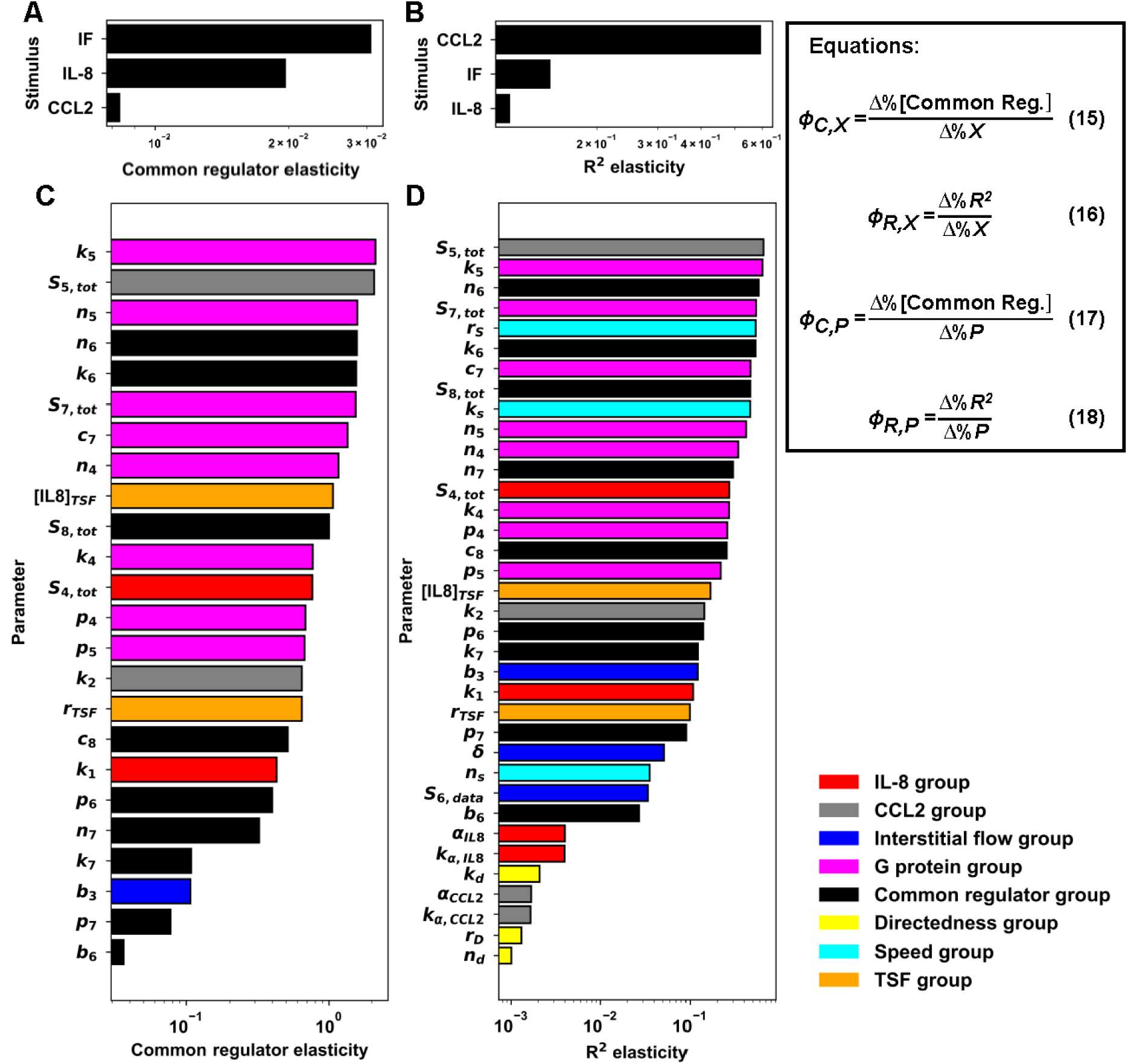
Sensitivity analysis that quantifies most influential stimuli and parameters on migration directedness and speed. The analysis considered (A) common regulator activation elasticity (Eq. 15) or (B) coefficient of determination (*R*^2^) elasticity (Eq. 16) in response to varying the magnitude of each extracellular stimulus. Also considered was (C) common regulator activation elasticity (Eq. 17) or (D) *R*^2^ elasticity (Eq. 18) in response to varying the magnitude of each parameter. The denominators Δ%*X* and Δ%*P* respectively represent a 20% change in the stimulus or the parameter under study. The numerator Δ%[Common Reg.] and Δ%*R*^2^ respectively represent the resulting percent change in concentration of active common regulator or the *R*^2^ value between the model predictions and the *in vitro* experimental data. All plots are base-10 log scale.

Then, a stimulus sensitivity analysis focusing on the *R*^2^ value was conducted to quantify the effect of varying the magnitude of each stimulus on the degree of agreement between the model and *in vitro* experimental data (Eq. 16) (Fig. 7b). This was done in order to determine the influence of each stimulus on the trends in directedness and speed across different stimuli as opposed to the concentration of active common regulator (which determines the magnitudes of directedness and speed). The influence of CCL2 on the *R*^2^ value was much greater than that of IF, which itself was greater than that of IL-8. This analysis suggests that while IF had the greatest influence on the concentration of active common regulator, CCL2 had the dominant influence on the trends in directedness and speed across different stimuli. Taken together, the considerable difference in the influence of the three modelled extracellular stimuli suggests that each plays a different role in the regulation of the downstream directedness and speed signals.

Focusing on the kinetics of individual reactions, a parameter sensitivity analysis was conducted to analyze how a change in a parameter affected the concentration of active common regulator (Eq. 17) (Fig. 7c). Additionally, parameters were classified according to their associated signaling protein or motif to assess the influence of reactions associated with each network element on the overall downstream effect. Results revealed that parameters associated with G protein activation and turnover were in general the most influential, with particular influence attributed to parameters governing CCL2-induced G protein activation (*k_5_* and *n_5_*). Other highly influential parameters included *S_5,tot_*, which dictates the total concentration of CCL2 receptor CCR2, and two parameters associated with common regulator activation in response to CCL2 signaling (*n_6_* and *k_6_*). Overall, this analysis suggests an important role for CCL2 signaling, in particular, and G-protein mediated signals, in general, in the regulation of the common regulator and thus migration directedness and speed.

A second parameter sensitivity analysis was conducted to quantify how varying the magnitude of each parameter changed the *R*^2^ value between the *in vitro* experimental data and model outputs (Eq. 18) (Fig. 7d). This analysis sought to determine the influence of each parameter on the trends in directedness and speed across different stimuli as opposed to the concentration of active common regulator. Again, parameters were classified according to their associated signaling protein or motif. The most influential parameter was *S_5,tot_*. Parameters associated with G protein activity were also among the most influential parameters, including *k_5_* (the second most influential parameter) which partially governs the rate of CCL2-induced G protein activation. Other highly influential parameters included those associated with the activation and turnover of the common regulator and the speed response to common regulator activity. Among the least influential parameters were those associated with the directedness response to common regulator activity. Overall, this analysis suggests that G protein signaling, particularly that induced by CCL2, also plays a dominant role in determining the trends across different stimuli and thus the agreement between *in vitro* experimental data and model-generated outputs.

## DISCUSSION

IF is an important tumor-associated biophysical factor that contributes toward cancer progression and poor patient survival [1,11–14]. Recent evidence suggests that IF also promotes the pro-tumor M2-polarization and migration activity of macrophages [51]. Moreover, clinical data demonstrate that macrophage density strongly correlates with increased metastasis [27]. Such findings suggest a plausible link between IF, macrophage activity (including their migration) and cancer metastasis, a research area that has not been widely studied [27]. In addition, macrophage migration is driven by tumor-secreted cytokines [36–40], which suggests that IF could act jointly with biochemical cues to affect macrophage migration. In this study, we demonstrate, for the first time, that IF can act in concert with tumor-secreted cytokines or factors (TSF) to regulate macrophage migration through a 3D *in vitro* TME-related ECM.

Previously, Li *et al*. observed that IF-exposed mouse macrophages migrated faster than non-treated control macrophages [51]. Following their work, we were interested to explore the results generated using a co-culture set-up of tumor cells and human primary macrophages, which more closely resembles the dynamic exchange between these cell types in the *in vivo* human TME. Using a two-gel channel set-up with tumor cells and macrophages in co-culture in adjacent but separate channels, we could delineate between the effects of TSF and tumor-originating IF on macrophage migration. We observed that IF-exposed macrophages migrated faster and with increased directedness than non-treated controls which agrees with Li *et al*.’s work. To our knowledge, our model is the first co-culture system that investigates the role of tumor IF on immune cell migration.

The Panc1 cell line was used to create the *in vitro* co-culture IF system. Panc1 originates from PDAC which clinically represents one of the most malignant of cancers with one of the highest death rates [78, 79]. PDAC metastasis correlates with a high macrophage infiltrate [55,80–84]. There are also numerous reports of cytokines underlying PDAC’s aggressive biology [85–88]. Therefore, PDAC seemed an appropriate cancer-immune model for investigating the effect of macrophage migration in response to PDAC-specific TSF, where the effect of IF can additionally be assessed. Also, our study integrated *in silico* and *in vitro* methods, an approach that has been highly effective in deriving insight into cell migration mechanisms [89–92]. Specifically, we demonstrated that by simulating *in vitro* conditions, an *in silico* signaling network model could be obtained associating key TSF (specifically IL-8 and CCL2), IF and macrophage migration. In turn, the *in silico* model was validated *in vitro* by adding different concentrations of exogenous IL-8/CCL2 or blocking antibodies against these cytokines and evaluating macrophage migration.

In our study, we first observed that the exposure of macrophages to IF or TSF induced a comparable increase in their migration directedness and speed. Interestingly, the non-additive effect of combining IF and TSF suggests that both IF and TSF could commonly regulate downstream macrophage migration when these stimuli are at saturating levels. Then, through a multiplex cytokine array of tumor-conditioned media, we identified IL-8 and CCL2 as the most probable cytokines driving the migration activity that we observed in the 3D *in vitro* system. In PDAC, acquisition of IL-8 and its receptors CXCR1 and CXCR2 on tumor cells [93, 94] and macrophages [95, 96] correlates with tumor invasion [97–99] and the metastatic potential of solid tumors in murine models [100–105] and patients [106, 107]. Similar to IL-8, the secretion of CCL2 and macrophage expression of its receptor CCR2 has been observed in the tumor tissues of patients with advanced metastasis [108–114]. Moreover, CCR2 blockade in a PDAC mouse model could deplete macrophages from the primary tumor to reduce metastasis [115]. Therefore, we focused on IL-8 and CCL2 in our study by virtue of their important role in regulating macrophage migration and PDAC metastasis. Notably, because IL-8/CCL2 are also implicated in the biology of other cancer types [42,116,117], our present findings can also be generalized to other cancers.

To understand the migration mechanism underlying our *in vitro* observations, we developed an *in silico* signaling network model to associate IL-8, CCL2, IF and macrophage migration. To develop this model, we first referenced key literature describing the intracellular signaling pathways associated with IL-8 and CCL2 [38–40], and studies concerning IF-induced cell migration mechanisms based on these cytokines [51,77,118]. Our findings show that IL-8 [66,68,69] and CCL2 [73, 119] activate signaling that results in the post-translational regulation of small monomeric GTPases of the Rho-family, leading to macrophage migration through the polymerization and retraction of the actin cytoskeleton. Specifically, in CCL2 signaling, extracellular chemokine CCL2 binds to and activates the chemokine receptor CCR2 expressed on the cell membranes of macrophages. Subsequently, the C-terminal intracellular domain of CCR2 activates intracellular signaling proteins, including phosphatidylinositol-3-kinase (PI3K), which eventually results in the activation of various Rho GTPases, in particular Rac, regulating cytoskeletal reorganization and cell migration [67,70–72,119]. In the case of IL-8 signaling, extracellular chemokine IL-8 binds to and activates receptor CXCR1/2, activating heterotrimeric small G-proteins, G(α,β,γ), which then promote the activation of Rho GTPases. Notably, other signaling activity downstream of CXCR1/2 and G(α,β,γ) also activates PI3K and FAK.

In addition, IF triggers a process known as outside-in signaling where it engages multiple extracellular signals to activate cell membrane-bound integrins that then initiate intracellular cytoplasmic signaling [118]. These extracellular signals include the binding of integrins to respective ligands in the ECM, and various mechanical forces originating from IF including fluid shear stresses. Specifically, each stimuli induces a conformational change in integrin, activating the cytoplasmic signaling element of the protein and allowing it to interact with other signaling molecules involved in intracellular signaling cascades [118]. Additionally, the activation of integrin activates FAK and Src, stimulating Rho GTPases which drive macrophage cytoskeletal reorganization and migration [51, 77]. Based on these findings, we hypothesized that IL-8, CCL2 and IF commonly regulate a group of Rho GTPases, including CDC42, Rac1 and/or RhoA [38–40], and this regulates macrophage migration. Additionally, the similar trends exhibited by both directedness and speed in response to all tested stimuli could be explained simply by a common regulator, as opposed to more complex interactions between largely independent signaling pathways. Finally, any attempt to fully model the highly complex signaling associated with migration regulation (that is downstream of this common signaling point) might greatly complicate the model with no added insight about how these signals are integrated. Thus, we depicted directedness and speed as phenomena indirectly induced by this common regulator, with intermediate signaling described by respective response functions that depend on the concentration of active common regulator.

To further validate our proposed model, we conducted a number of simulations and model analyses. We first demonstrated that the model was capable of reproducing the same trends of directedness and speed that we observed *in vitro* in response to the exposure to various combinations of IL-8, CCL2 and IF. Quantifying this agreement by calculating the *R*^2^ value across all modelled directedness and speed data, we determined that the model successfully accounts for over 83% of the variation in the experimental data. Additional Student’s *t*-tests further confirm the reproducibility of the model as there was no substantial difference between the model’s predicted values and *in vitro* measurements at a point-by-point level. In line with *in vitro* experimentally observed macrophage migration behaviors, our model also displayed an OR gate-like behavior, where the exposure to a single stimulus (of biologically consistent magnitude) results in maximum common regulator activation. We highlight that such behavior is essential for integrating multiple inputs to produce a stable output regardless of input number. It is also an important, though conditional, example of redundant signaling where multiple signaling pathways lead to the same downstream effect.

Although TSF contains cytokines in addition to IL-8 and CCL2, the connections between the IL-8, CCL2 and IF-induced, integrin-β2-mediated, signaling appear to suitably explain the *in vitro* experimentally-observed macrophage migration. The relatively dominant role of IL-8 and CCL2 is reflected in a second set of experimental data where *in vitro* antibody blockade of IL-8 and/or CCL2 substantially inhibited the TSF-mediated increase in migration. Moreover, the simultaneous exposure of macrophages to sub-optimal concentrations of both IL-8 and CCL2 was able to achieve a similar effect on migration directedness and speed as their exposure to only TSF. Of note, we recognize that IF and TSF do not appear to act additively in the tumor cell-macrophage co-culture set-up where cytokines are likely to be at saturating levels (confirmed by the similar increase in directedness/speed between the TSF condition and concentration of 100 ng/mL that was used to intentionally induce saturation, Supplementary Fig. S2). Instead, the increase in directedness/speed with either (1) a combination of IL-8 and CCL2 with each cytokine at a sub-optimal level (25 ng/mL) or (2) sub-optimal level of either cytokine with IF suggests that synergies are possible at non-saturating cytokine concentrations. Such synergies have been previously demonstrated experimentally for macrophages, and have been determined to depend on the unimpaired activity of both CXCR1/2 and CCR2 receptors, suggesting the intracellular integration of both cytokine signals as the primary synergistic mechanism [75].

A comparison of model predictions and *in vitro* data of these antibody blockade and sub-optimal concentration experiments further validates our network architecture and kinetics assumptions. First, consistent with experiments, the behavior with saturating cytokine concentrations was not observed in response to diminished concentrations. Furthermore, by accurately capturing the signaling instigated by intermediate and sub-optimal cytokine concentrations, our model showed that it is not a simple all-or-nothing OR-gate, but a nuanced model that can capture the response across a gradient of stimuli. Second, the model accurately reproduced migration behaviors associated with the antibody blockade of IL-8 and/or CCL2, reinforcing its capability to accurately reproduce experimentally relevant phenomena and capture intermediate signaling with diminished cytokine concentrations (as quantified by the *R*^2^ values between modelled and *in vitro* experimental data).

We then conducted a number of sensitivity analyses to determine the most influential aspects of the network on the network response magnitude and the data trends. Stimulus sensitivity analysis revealed that IF was more influential than either cytokine on the concentration of active common regulator, whereas CCL2 was the most influential extracellular stimulus on the directedness and speed trends between various extracellular stimuli (as quantified by the *R*^2^ values between modelled and *in vitro* experimental data). Parameter sensitivity analysis then revealed that reactions associated with G protein activation and CCL2 signaling were the most influential on the resulting concentration of active common regulator as well as the trends between various extracellular stimuli (quantified by the *R*^2^ values between modelled and *in vitro* experimental data). Also, the activation and turnover of the common regulator itself was important to both active common regulator magnitude and data trends. Notably, although the magnitude of IF velocity has a significant influence on the concentration of active common regulator, the parameters associated with IF signaling were not among the most influential parameters on the concentration of active common regulator.

Importantly, our findings substantiate the idea that IF contributes to cancer invasiveness through enhancing macrophage migration. Indeed, as macrophages would have a heightened capacity to migrate through the 3D ECM, there would be increased likelihood for them to interact with and hence support cancer cell migration in the process of metastasis. The supportive function of macrophages toward cancer metastasis was previously demonstrated, where media from tumor-conditioned macrophages increased the expression of EMT genes in a low EMT-score tumor cell line [55]. Moreover, media conditioned from IF-exposed macrophages could increase the speed of cancer cell migration through a 3D matrix [51], suggesting that IF could support the capability of macrophages to promote tumor cell invasion. These findings support the view that macrophages play a pivotal intermediary role between the stimulus of IF and the output of cancer cell invasion through 3D ECM. Notably, other stromal cells in the TME may also respond to IF and further influence macrophage and/or cancer cell migration. For example, cancer associated fibroblasts (CAFs) can secrete ECM to remodel the TME matrix and this can influence cell migration in the TME [120]. Future studies could therefore incorporate other TME-related cells, such as CAFs, to evaluate their contribution to the relationship between IF and the metastasis process.

Therefore, our work presents an integrated *in silico*-3D *in vitro* approach to evaluate the effect of IF and TSF on macrophage migration. Here, we developed a signaling network model identifying key stimuli and intermediary proteins that drive macrophage migration, thus identifying potential therapeutic targets for inhibiting macrophage migration (which evidently associates with their capability to support cancer cell invasion). Our model was established using cancer cell lines and macrophages derived from the *in vitro* differentiation of blood-isolated monocytes. By incorporating patient-derived tumor explants and autologous macrophages, our platform could potentially facilitate high-throughput preclinical screening of therapies for personalized treatment. Our work also forms the basis for developing a companion diagnostic that comes with a biophysical component such as IF for identifying patient responders. In addition, as IF presents a physical barrier to the effective penetration of drugs into deeper regions of the TME, our model could be used to screen and guide the design of therapeutics to optimize their transport efficiency. Importantly, this work contributes toward an improved understanding of the signaling mechanism associating IF, macrophage motility and cancer metastasis, an area that should be studied more extensively to improve cancer treatment.

## Supplementary Material

Supplementary material is available on the bioRxiv server.

## Author Contributions

S.L., W.S.C., G.A. and R.K. designed the study and *in vitro* experiments. S.L., G.A. and A.T. conducted *in vitro* experiments. R.S., F.L. and F.S. developed the computational model. R.S. implemented the model and conducted simulations. D.K. transfected the human cell lines used in the study. S.L. and R.S. analyzed the results. W.S.C., G.A., M.Z. and R.K. supervised the study and acquired funding. All authors interpreted the results, reviewed and edited the manuscript.

## Acknowledgements

The authors would like to acknowledge the SIgN Multiplex Analysis of proteins (MAP) platform for performing the multiplex cytokine array and R&D systems for performing the protein array of cell culture supernatant. We also thank Vicnesvari T. (Singapore-MIT Alliance for Research and Technology, SMART) for assisting with fabricating microfluidic devices, Giovanni S. Offeddu (Massachusetts Institute of Technology) and Luca Possenti (Politecnico di Milano) for their guidance on performing FRAP analysis and Ran Li (Massachusetts General Hospital) for providing scientific and technical input.

