## Supplementary for "Integrated *in silico* and 3D *in vitro* model of macrophage migration in response to physical and chemical factors in the tumor microenvironment"

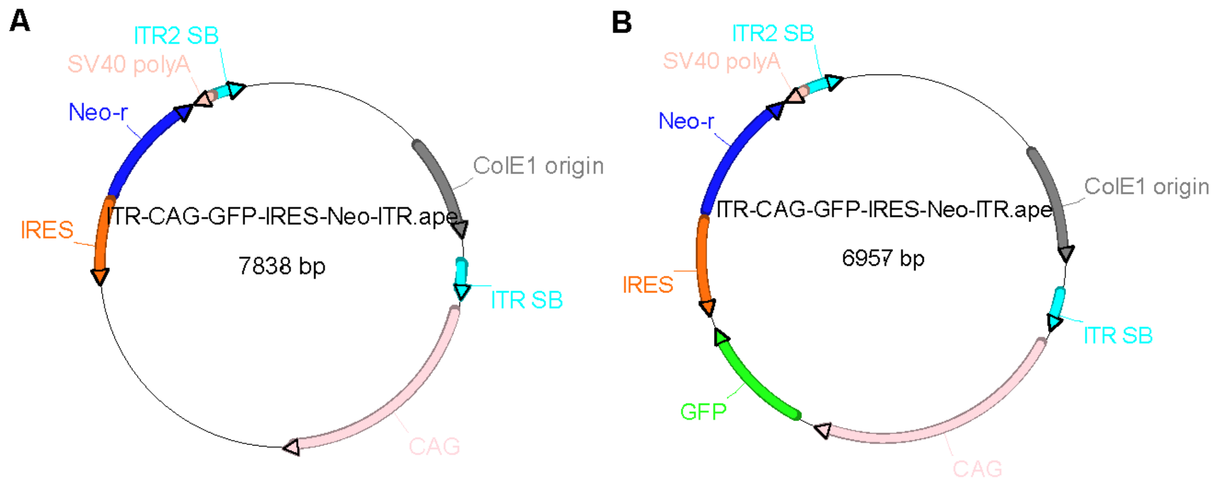


**Supplementary Figure S1.** (A) ITR-CAG-DEST-IRES-Neomycin-ITR (control plasmid) or (B) ITR-CAG-GFP-IRES-Neomycin-ITR (GFP plasmid) used to generate the GFP stable cell lines.

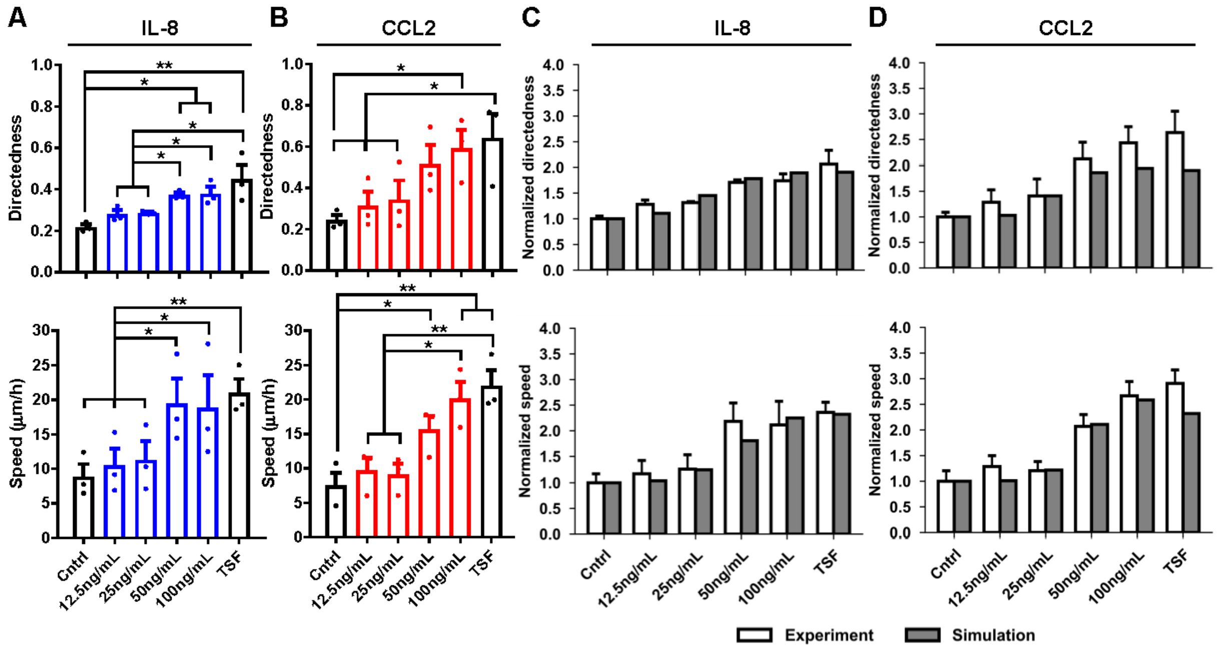


**Supplementary Figure S2.** Macrophage migration directedness (top panel) and speed (bottom panel) in response to different concentrations of (A) IL-8 or (B) CCL2 that were added to a macrophage monoculture in the absence of interstitial flow. Non-significant difference between experimental (white) and model (grey) predictions for normalized directedness and speed for titration data of (C) IL-8 or (D) CCL2. Data are shown as the mean ± SEM (n = 3), where statistical significance was determined using a (A, B) one-way ANOVA with Holm-Sidak's multiple comparisons test with * $P \leq 0.05$ and ** $P \leq 0.01$, or (C, D) Student's *t*-tests that compared between *in vitro* experimental data and model-generated data at a pointwise level with $P \leq 0.05$. (*Cntrl*: control, *TSF*: tumor-secreted factors)

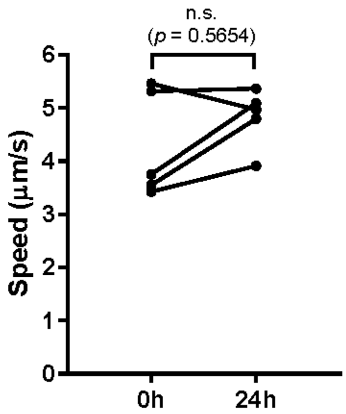


**Supplementary Figure S3.** Fluorescence recovery after photobleaching (FRAP) analysis of flow velocities within the gel of the 3D *in vitro* microfluidic model of interstitial flow (IF). Mean flow velocities at the start and end of the 24 h of IF treatment are shown. Velocities fall within 3-5 µm/s which correspond with physiological values of tumour IF. Data are shown as the mean (n = 4), where statistical significance was determined using a Student's *t*-test. (*n.s.*: not significant)
